# Common transcriptional programme of liver fibrosis in mouse genetic models and humans

**DOI:** 10.1101/2020.11.02.364901

**Authors:** Kaja Blagotinšek Cokan, Žiga Urlep, Miha Moškon, Miha Mraz, Xiang Y. Kong, Winnie Eskild, Damjana Rozman, Peter Juvan, Tadeja Režen

## Abstract

Multifactorial metabolic diseases, such as non-alcoholic fatty liver disease, are a major burden of modern societies and frequently present with no clearly defined molecular biomarkers. Herein we used systems medicine approaches to decipher signatures of liver fibrosis in mouse models with malfunction in genes from unrelated biological pathways. Enrichment analyses of KEGG, Reactome and TRANSFAC databases complemented with genome-scale metabolic modelling revealed fibrotic signatures highly similar to liver pathologies in humans. The diverse genetic models of liver fibrosis exposed a common transcriptional programme with activated ERα signalling, and a network of interactions between regulators of lipid metabolism and transcription factors from cancer pathways and immune system. The novel hallmarks of fibrosis are downregulated lipid pathways, including fatty acid, bile acid, and steroid hormone metabolism. Moreover, distinct metabolic subtypes of liver fibrosis were proposed, supported by unique enrichment of transcription factors based on the type of insult, disease stage, or sex.

## Introduction

Fibrosis is a common companion of skin, lung, kidney and liver diseases, while it can affect virtually every organ. It is characterised by excessive deposition of connective tissue components, which leads to tissue remodelling and organ malfunction. High mortality is associated with fibrotic diseases. The progress in development of anti-fibrotic drugs is slow, especially for individual fibrotic diseases where mechanisms beyond are not clear. There is a need to unravel the »core« fibrotic pathways across different fibrotic diseases as well as across the same type of fibrotic disease that can arise from a multitude of causes. It is believed that in addition to common fibrotic programmes, other factors influencing fibrotic disease susceptibility may be distinct, with disease-specific and organ-specific risk factors (Distler et al. 2019).

Liver fibrosis is a characteristic of the progressive liver pathologies defined by accumulation of collagen, smooth-muscle actin, hydroxyproline, etc., and is one of the hallmarks of the advanced stages of non-alcoholic fatty liver disease (NAFLD); currently called metabolism-associated fatty liver disease (MAFLD). This is a multifactorial disease with variable aetiology and no clearly defined molecular biomarkers for diagnosis, prognosis or progression (Eslam, Sanyal, and George 2020). The advanced disease stages include non-alcoholic steatohepatitis (NASH) and cirrhosis, both potentially leading towards liver cancer. The prevalence and hence the burden of the disease is increasing because of the lack of approved pharmacotherapies and low impact of prevention strategies (Younossi et al. 2018). In animal models, liver fibrosis is a result of a chronic liver injury induced by different factors, which range from alcohol, diets, toxins, drugs, bile duct ligation, genetic modifications, and others. Each type of liver injury activates a specific programme at cellular and molecular levels (Liedtke et al. 2013) and if sustained, disease progresses to further stages (Sircana et al. 2019).

In humans, NAFLD and NASH have many faces as clinical manifestations are highly heterogeneous (Friedman et al. 2018). Cirrhosis may or may not be present, not all patients show abnormal blood parameters, comorbidities, such as diabetes and obesity, vary, the presence of fibrosis and steatosis is not uniform. Clinical drug trials consistently show that targeting NAFLD histological features does not always result in disease resolution. For example, reduction of steatosis did not improve other histological outcomes of NAFLD (Bril et al. 2020) and elevated steatosis did not always associate with worsening of fibrosis (Rinella et al. 2019). All this indicates that there are potentially different subtypes of NAFLD patients. In concordance, a recent study identified three NAFLD subtypes in relation to methionine/folate cycle according to serum metabolite signature, also predicting the progression to NASH (Alonso et al. 2017). The latest recommendation was subcategorization of NASH patients to identify those who will be best suited for specific treatments in clinical trials (Rinella et al. 2019).

As resolution of fibrosis is one of the endpoints of clinical trials, we can benefit from a variety of mouse models that develop progressive fibrosis similarly as humans. In our previous work we discovered that hepatocyte-specific *Cyp51* (cytochrome P450 or lanosterol 14α-demethylase) knockout (LKO) males and females develop liver fibrosis (without steatosis or cholestasis) due to the block of cholesterol synthesis (Lorbek et al. 2015). Among metabolic alterations were deregulated sterol intermediates, decrease in hepatic cholesterol and its esters, modified bile acid composition and elevated plasma total cholesterol and HDL in a sex-specific manner. Similarly, the whole-body knockout (KO) of *Glmp* (glycosylated lysosomal membrane protein) presented with liver fibrosis although the function of this lysosomal protein remains to be clarified (Kong et al. 2014). Among metabolic alterations in *Glmp* KO mice are increased liver bile acids and infiltration of inflammatory cells (Nesset et al. 2016). Decreased blood glucose, triglycerides (TAG) and non-esterified fatty acids were also observed, together with increased liver TAGs although liver steatosis was not confirmed histologically (Kong et al. 2015).

If two such different KO models both result in a similar liver phenotype, we hypothesized that it might be possible to determine a common fibrotic signature from multiple mouse models. We focused on single gene knockouts that develop histologically confirmed liver fibrosis (with or without steatosis or cholestasis) without additional dietary or chemical insults, preferably in both sexes, and with well annotated transcriptome data. In addition to *Cyp51* LKO and *Glmp* KO, the fibrotic phenotype also develops in the liver knockout of Notch signalling pathway repressor *Rbpj* (recombination signal binding protein for immunoglobulin kappa J region) due to impaired bile duct maturation causing obstructive bile acid flow, accumulation of bile acids, necrosis, and severe cholestasis with progression to hepatocellular carcinoma (HCC) (Tharehalli et al. 2018). Fibrosis is also a feature of the hepatocyte-specific *Ikbkg* (*Nemo*, Inhibitor of kappaB kinase gamma) knockout males with steatohepatitis and HCC through changing the response to inflammation (Beraza et al. 2009). Increased oxidative stress was present, mitochondria appeared to be affected, irregular glycogen deposits were observed, and serum glucose was decreased while serum TAG and cholesterol levels were unchanged (Luedde et al. 2007). We aimed to study the fibrotic signatures of both sexes; however, only the *Cyp51* LKO transcriptome data include also females. Using functional comparative analysis of gene expression based on KEGG, Reactome and TRANSFAC databases, as well as genome-scale metabolic models (GEMs), we could identify multiple common fibrotic transcriptome signatures, with high similarity to human NAFLD and NASH. The hallmark is downregulation of metabolic pathways and upregulation of immune system-related pathways and pathways in cancer. We provide also new insights into the function of GLMP in the liver and propose “universal” fibrosis-related biomarkers with a touch on sex dependencies.

## Results

### Similar transcriptome alterations caused by different genetic defects

The characteristics of genetic mouse models are summarized in **Supplementary Table 1**. They were all adult males with C57BL/6J genetic background, in addition to a group of *Cyp51* LKO females, with histologically confirmed fibrosis, increased inflammation (except *Rbpj* LKO where it was not measured), and final progression to liver tumours or dysplastic nodules. The *Cyp51* LKO and *Glmp* KO did not present with histological steatosis and cholestasis. We analysed the liver transcriptome data and compared the differentially expressed genes (DEG) between these genetic models of fibrosis (**Suppl. Table 2**). At the intersection we observed a higher number of upregulated (59) than downregulated genes (3) (**Fig. 1**). A higher percentage of upregulated genes was common also when we compared the overlap of DEG between each pair of fibrotic models (**Suppl. Table 3**). From the common DEG, the majority has a function at the plasma membrane or extracellular space or are involved immune response pathways, among them *Tgfbr2, Tgfbi* from TGF-β signalling, and the lipoprotein lipase (**Fig. 2A**). TGFBI was recently confirmed to be strongly associated with NAFLD and cirrhosis in humans (Niu et al. 2019). These results lead us to a conclusion that the common transcriptional programme of liver fibrosis is largely a part of the upregulation of genes, while downregulation of genes is more associated with the unique programme probably related to the type of the insult, disease stage or sex.

**Figure 1.**
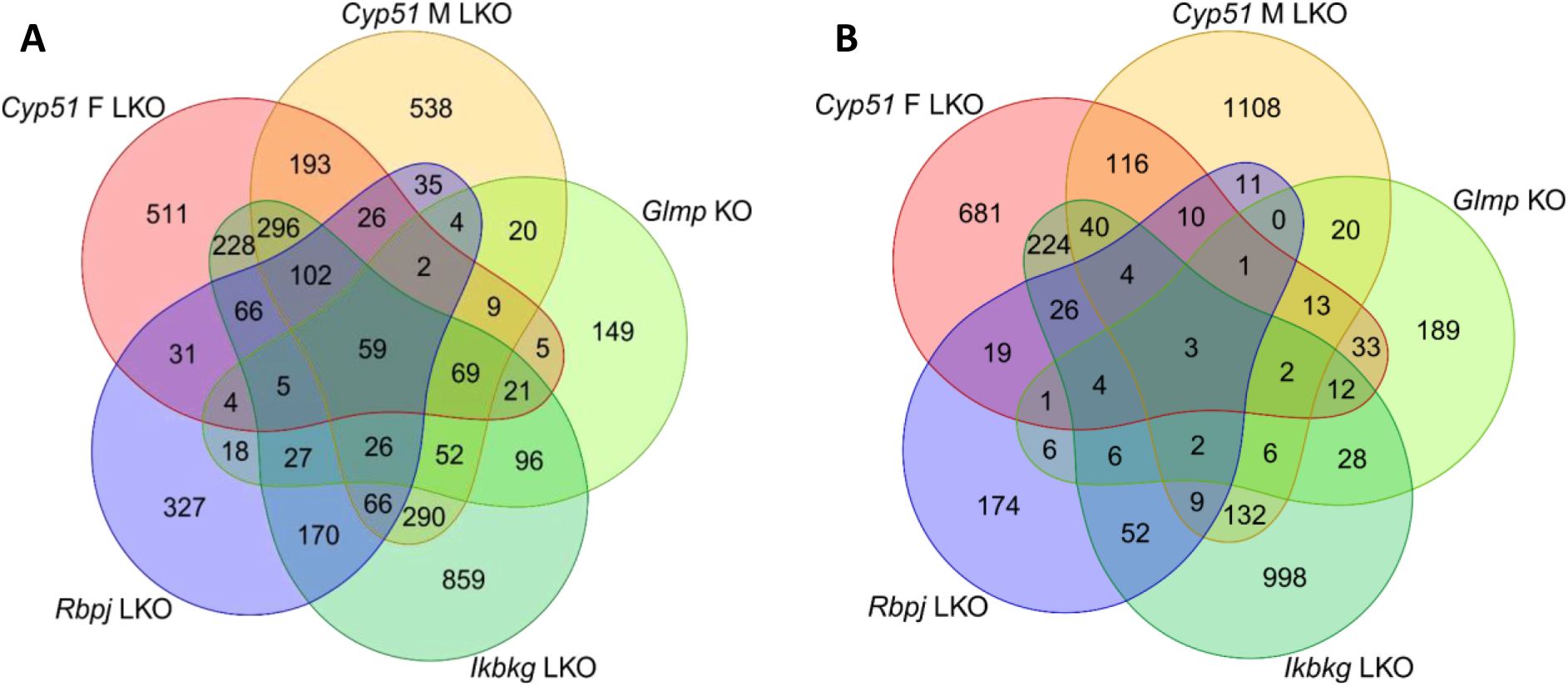
Venn diagrams of DEGs in mouse genetic models of liver fibrosis. **A.** Upregulated DEGs. **B.** Downregulated DEGs.

**Figure 2.**
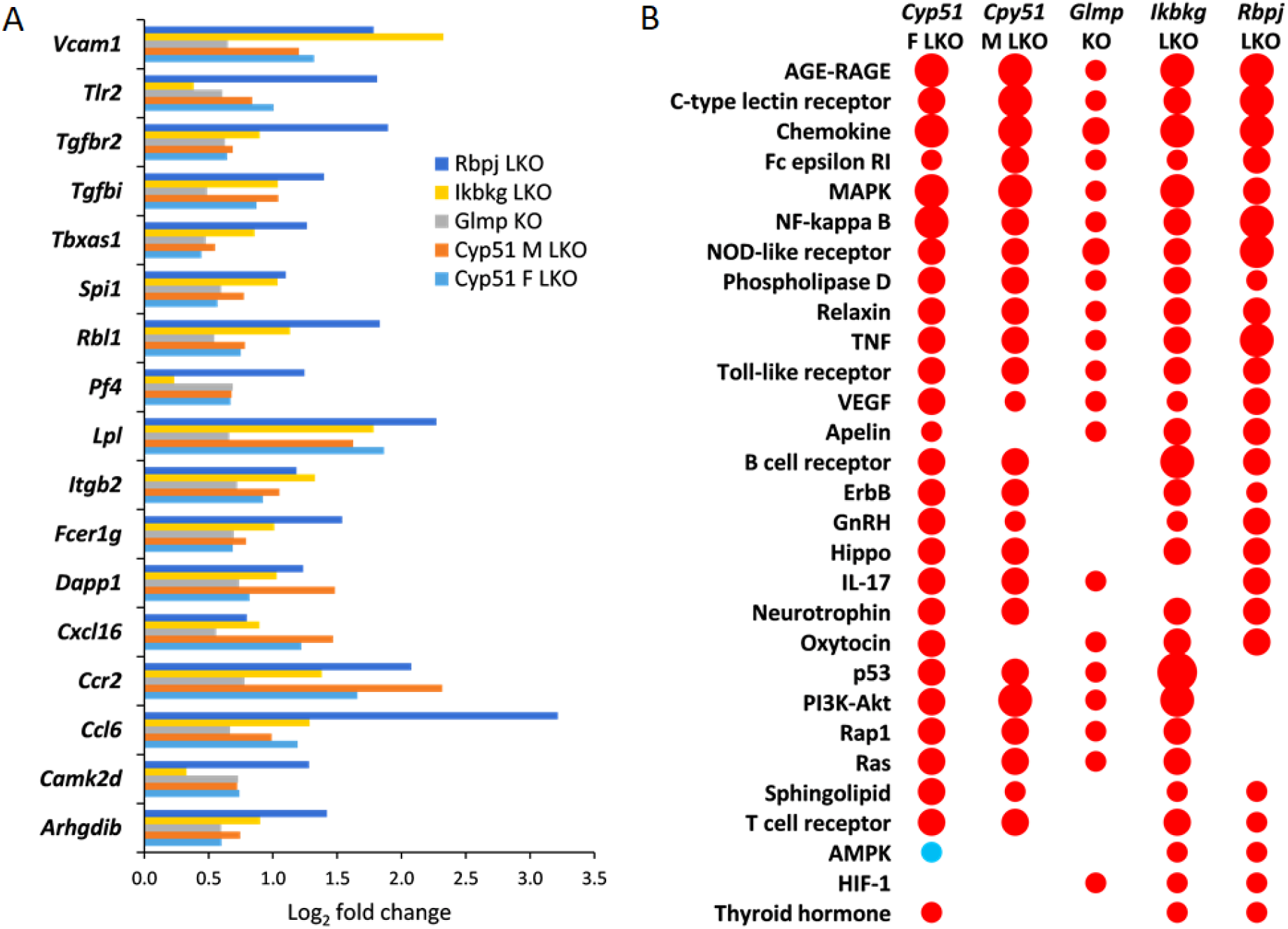
Common significantly enriched KEGG signalling pathways and DEGs. **A.** Log_2_ fold change in expression of selected common DEGs in all fibrotic models. Only DEGs, which are a part of common enriched signalling or metabolic KEGG pathways are presented. Log_2_ fold change represents Log_2_ ratio between average KO *vs* WT. **B.** KEGG signalling pathways common to at least three mouse genetic models of liver fibrosis as calculated using pGSEA are presented. Red indicates positive enrichment and blue negative. The size of the bubble reflects the fold change of each pathway.

### Common KEGG and Reactome pathways in different fibrotic models

Positively enriched KEGG (65) and Reactome (52) pathways common to all liver fibrotic models have roles in response to liver injury, repair, haemostasis, cancer, regulation of metabolism, development of fibrosis, and activation of the immune system (**Suppl. Table 4 and 5**). Both pathway analyses confirmed that models are most similar in positively enriched pathways (**Suppl. Table 3**). At the intersection of different genetic models, there were many positively enriched KEGG pathways indicating an activation of a common transcriptional program in fibrosis regardless of the type of injury, age or sex (**Fig. 2B**). Common enriched Reactome and KEGG pathways were, with few exceptions, always enriched in the same direction in the models. An exception was, for example, AMPK signalling, which was positively enriched in *Ikbkg* LKO and *Rbpj* LKO, but negatively in female *Cyp51* LKO (**Fig. 2B**). Pathway analysis revealed that cellular organelles are affected during progression of fibrosis. For example, peroxisome pathways (*Peroxisomal protein import, Peroxisome* pathway) and *Autophagy* were negatively enriched while *Lysosome* was enriched positively. Interestingly, with exception of *Glmp* KO and the male *Cyp51* LKO, *HCC pathway* was enriched positively, indicating an early commitment of liver cells towards cancer in these models.

### Mouse genetic models show overlapping transcriptome signatures with human NAFLD and NASH

It is important to compare the mouse genetic models of hepatic fibrosis to human NAFLD and NASH. We selected a list of enriched KEGG pathways calculated by GSEA in patients with NAFLD (N=27) and NASH (N=25) as presented in Teufel *et al*. (Teufel et al. 2016) where liver transcriptome data from NAFLD and NASH patients of both sexes were compared to control (N=39) and healthy obese (N=25) patients and also to mouse female diet models. In our analyses, we used the Teufel data for direct comparison of our pGSEA analyses from genetic models with data from patients with NAFLD and NASH (**Suppl. Table 6**). There were many common positively enriched KEGG signalling pathways between mouse genetic model and human NAFLD and NASH, which is in line with a common fibrotic programme (**Table 1**). Interestingly, the genetic fibrotic models had a much higher overlap of enriched KEGG pathways with human NAFLD and NASH compared to dietary mouse models reported in Teufel et al (Teufel et al. 2016). This was surprising since in humans, diet is supposed to be a crucial factor in majority of NAFLD cases. However, we cannot exclude that some of the observed distinctions were not due to differences between humans and mice or due to diet, but might be linked to sex. Our mouse models were of both sexes while Teufel data included NAFLD and NASH patients of both sexes and only female dietary mouse models.

**Table 1:** Top enriched KEGG pathways common between genetic mouse models of liver fibrosis and human NAFLD and NASH from Teufel *et al* (Teufel et al. 2016). Presented are log_2_ fold changes comparing KO *vs* WT for mouse models and direction of enrichment for human NAFLD and NASH data, where + means enrichment and ns - not significant.

| KEGG pathway | NAFLD | NASH | *Cyp51* F LKO | *Cyp51* M LKO | *Glmp* KO | *Ikbkg* LKO | *Rbpj* LKO |
| --- | --- | --- | --- | --- | --- | --- | --- |
| Antigen processing and presentation | + | + | 1.00 | ns | 0.84 | 1.05 | ns |
| B cell receptor signalling pathway | + | + | 0.89 | 0.77 | ns | 1.06 | 0.91 |
| Cell adhesion molecules (CAMs) | + | + | 1.43 | 1.12 | 1.02 | 1.34 | 0.60 |
| Cell cycle | ns | + | 1.25 | ns | 0.50 | 2.40 | 1.40 |
| Chemokine signalling pathway | ns | + | 1.06 | 0.78 | 0.89 | 1.17 | 1.39 |
| Colorectal cancer | ns | + | 0.87 | 0.73 | ns | 0.84 | 0.47 |
| Cytokine-cytokine receptor interaction | ns | + | 0.76 | 0.77 | 0.71 | 0.93 | 1.21 |
| DNA replication | + | + | 0.99 | ns | ns | 1.01 | 1.23 |
| ECM-receptor interaction | + | + | 1.67 | 1.87 | 0.54 | 1.50 | 0.50 |
| Endocytosis | ns | + | 1.20 | 0.72 | ns | 0.93 | 0.65 |
| ErbB signalling pathway | ns | + | 0.80 | 0.56 | ns | 0.78 | 0.49 |
| Fc epsilon RI signalling pathway | + | + | 0.67 | 0.40 | ns | 0.50 | 0.75 |
| Fc gamma R-mediated phagocytosis | + | + | 1.08 | 0.93 | ns | 1.06 | 0.97 |
| Focal adhesion | + | + | 1.96 | 1.85 | ns | 1.65 | 0.67 |
| Hematopoietic cell lineage | + | + | 1.20 | 1.59 | 1.23 | 1.05 | 0.95 |
| Leukocyte transendothelial migration | + | + | 1.52 | 1.11 | 0.70 | 1.08 | 0.90 |
| MAPK signalling pathway | ns | + | 1.09 | 1.08 | ns | 1.26 | 0.70 |
| Natural killer cell mediated cytotoxicity | ns | + | 0.95 | ns | 0.54 | 0.87 | 0.80 |
| Neurotrophin signalling pathway | ns | + | 0.98 | 0.53 | ns | 0.63 | 0.60 |
| Pancreatic cancer | + | + | 1.01 | 0.76 | ns | 0.95 | 0.42 |
| Pathways in cancer | ns | + | 1.42 | 1.18 | ns | 1.31 | 1.21 |
| Phagosome | + | + | 1.71 | 1.48 | 1.10 | 1.70 | 1.53 |
| Regulation of actin cytoskeleton | ns | + | 1.77 | 1.39 | ns | 1.27 | 0.63 |
| Small cell lung cancer | + | + | 1.21 | 1.18 | 0.44 | 1.02 | 0.64 |
| T cell receptor signalling pathway | ns | + | 0.73 | 0.52 | ns | 0.63 | 0.40 |
| Toll-like receptor signalling pathway | ns | + | 0.85 | 0.70 | ns | 0.69 | 0.65 |
| VEGF signalling pathway | + | + | 0.48 | 0.28 | ns | 0.49 | 0.54 |

### Negative enrichment of metabolic pathways in fibrosis

We discovered that negative enrichment of metabolic pathways is a hallmark of fibrosis. From basic metabolism, bile and fatty acids (linoleic), steroid hormone, ketone, butanoate, nitrogen, heme and branched-chain amino acid-related KEGG and Reactome pathways were enriched negatively in the four genetic fibrotic models (**Table 2, Suppl. Table 4 and 5**). Most importantly, the upstream pathways regulating metabolism were also negatively enriched, such as several nuclear receptors, IGF1R and insulin signalling (**Suppl. Table 5**). In contrast, *Glycosphingolipid* and *Sphingolipid metabolism* and its signalling were positively enriched in all models except *Glmp* KO. These results indicate that the modulation of metabolic pathways happens regardless of the type of injury, metabolic state or sex and also in absence of dietary manipulation.

**Table 2:**
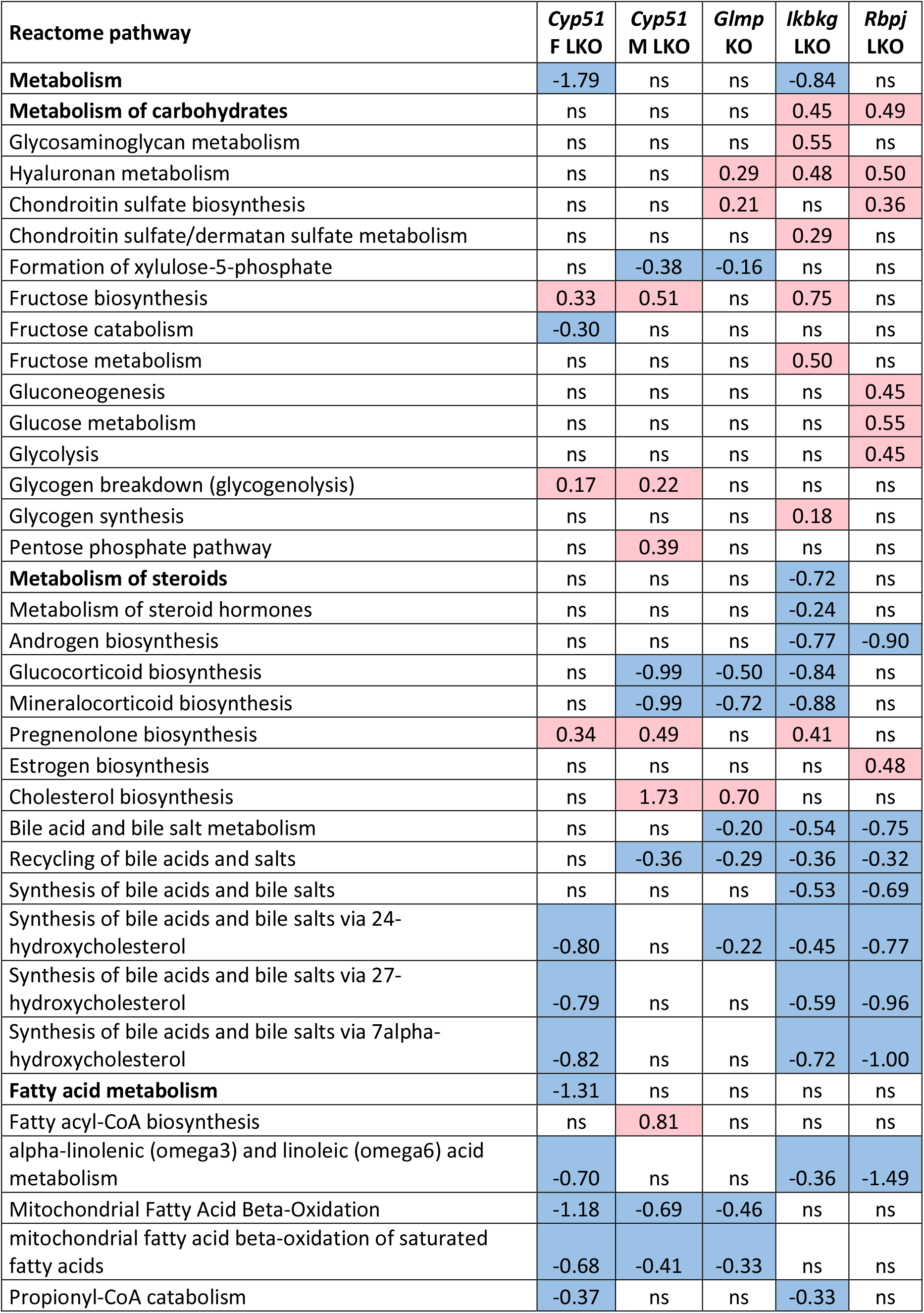
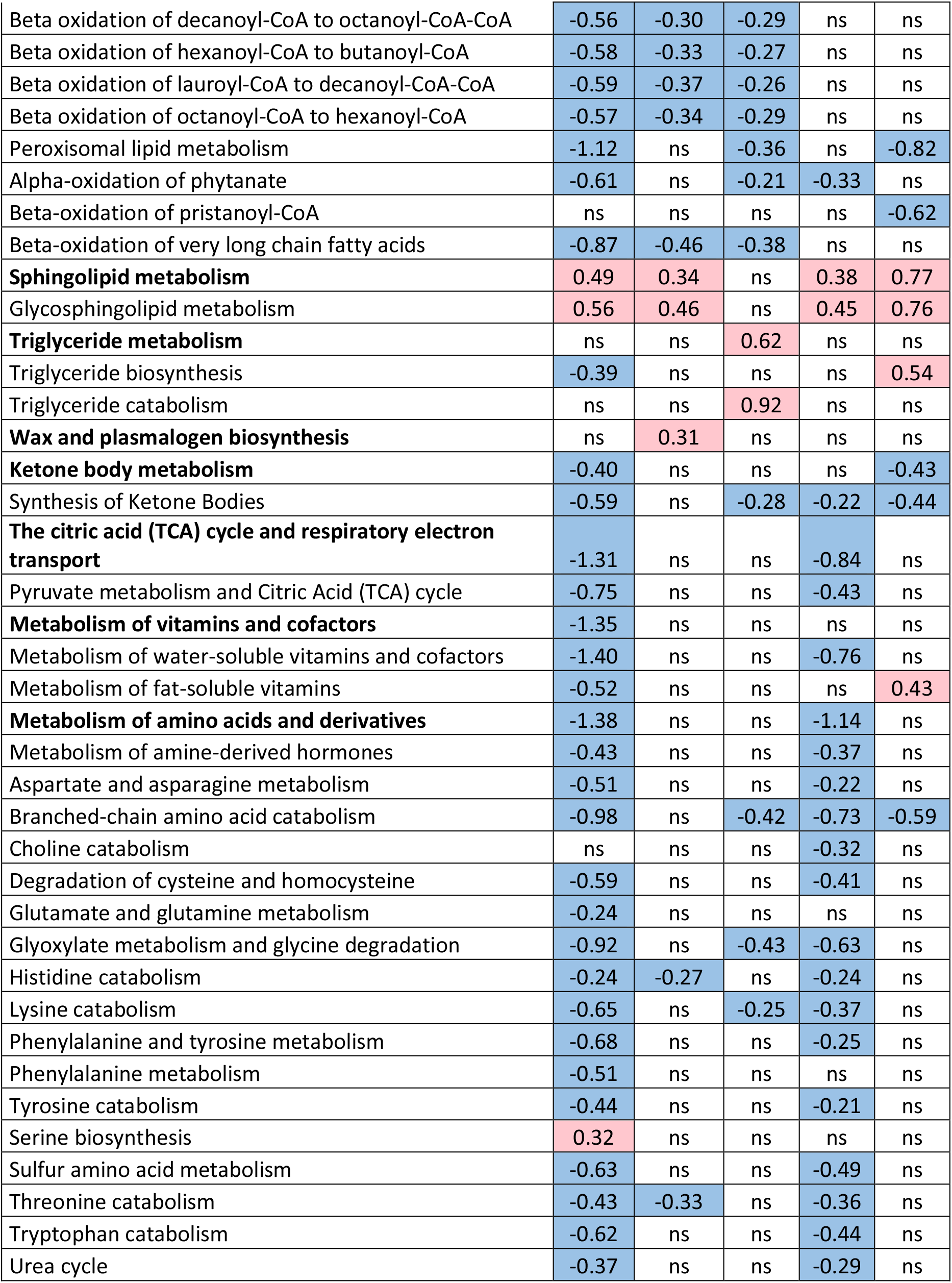
Selected statistically significantly enriched Reactome pathways from metabolism. Presented are log_2_ fold changes KO vs WT, red are upregulated and blue downregulated. ns – non-significant.

Focusing on more unique pathways and DEGs that are not common among models we observed distinct changes in metabolism of almost all amino acids, carbohydrates, vitamins, cofactors and energy metabolism, exposing female *Cyp51* LKO and male *Ikbkg* LKO models as the most affected (**Table 2**). These results propose specific metabolic programmes and existence of different metabolic subtypes depending on the type of injury, stage of fibrosis or sex. In conclusion, data from genetic fibrotic models exposed a wide array of metabolic rearrangements as the hallmark of the fibrosis program.

### Genome-scale metabolic models confirmed rearrangements in lipid metabolism pathways

Since metabolic rearrangements were enriched in pathway analyses, we used GEMs to simulate and predict metabolic fluxes at systems-level using transcriptome data. **Fig. 3** represents statistically significant changes in GEMs common to at least three models, with majority involved in fatty acids metabolism (synthesis, oxidation, transport) in different cellular compartments (cytosol, mitochondria, peroxisome). Importantly, several sterol related GEMs were affected by fibrosis, such as cholesterol and bile acid synthesis, cholesterol esters and steroid metabolism (**Fig. 3**). The levels of liver cholesterol, cholesterol esters and bile acids levels were decreased in *Cyp51* M LKO model, indicating that similar conditions might be present in other genetic models (Lorbek et al. 2015). Several other lipids were rearranged during fibrosis, such as glycerolipids, glycerophospholipids, sphingolipids, etc. GEM analyses also detected changes in carnitine shuttle, transport reactions and retinol metabolism. Again, each model exhibited a unique combination of metabolic rearrangement. Importantly, the GEM analyses confirmed the global rearrangement of lipid homeostasis as a part of the common programme in liver fibrosis. Moreover, GEM analyses indicated that the type of injury defined which cellular compartment was affected and in what way.

**Figure 3.**
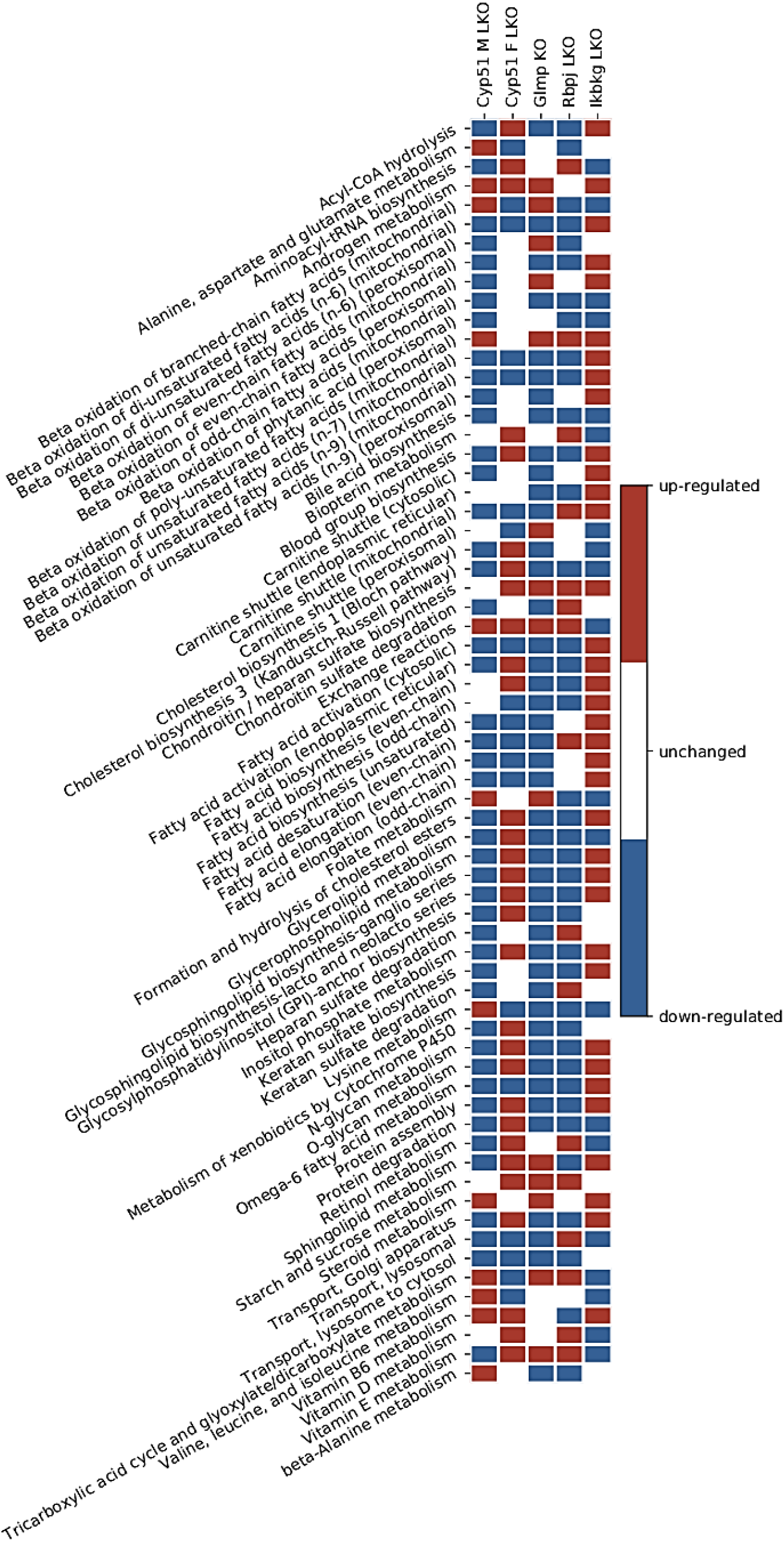
Statistically significantly perturbed GEM subsystems. Presented are subsystems common to at least three models of liver fibrosis

### Enrichment of transcription factors exposed variability in metabolic regulators

Transcription factors are upstream regulators of metabolism and are thus superior to metabolic pathways in KEGG and Reactome, adding another level of understanding. Majority of TFs were positively enriched (**Suppl. Table 3 and 7**). Twenty TFs were enriched in all genetic fibrotic models, among them ERα, NF-Y, c-ETS-1, etc., and additional 16 were common to four (**Suppl. Table 7**). All were enriched positively and involved in the regulation of immune system, cancer, metabolic or hormone pathways. Several nuclear receptors regulating lipid metabolism and estrogen receptor α were enriched, coinciding with enrichment of Reactome pathways related to estrogen receptor and nuclear receptors (**Fig. 4A**). Cytoscape analysis using STRING database revealed a network of interactions between the common TFs regulating lipid metabolism and common TFs regulating cancer pathways and immune system (**Fig. 4B**). It is important to note that the enrichment of TFs regulating lipid metabolism varied among the studied genetic models (**Fig. 4A**). For example, E2F-1, a mediator of sustained lipogenesis and contributor to hepatic steatosis, was enriched positively in *Ikbkg* LKO and *Rbpj* LKO, and negatively in *Glmp* KO, coinciding with histological findings. PPARγ:RXRα, RXRα and VDR were positively enriched in at least three models, LXRα and PPARα in two to three, while FXR was not enriched at all. Another known lipid regulator is SREBP1, which was positively enriched in *Rbpj* LKO, and negatively in *Ikbkg* LKO. These data indicate that the combination of enriched TFs, the regulators of metabolism, could depend on genetic background and could be used to predict the metabolic subtypes of fibrosis.

**Figure 4.**
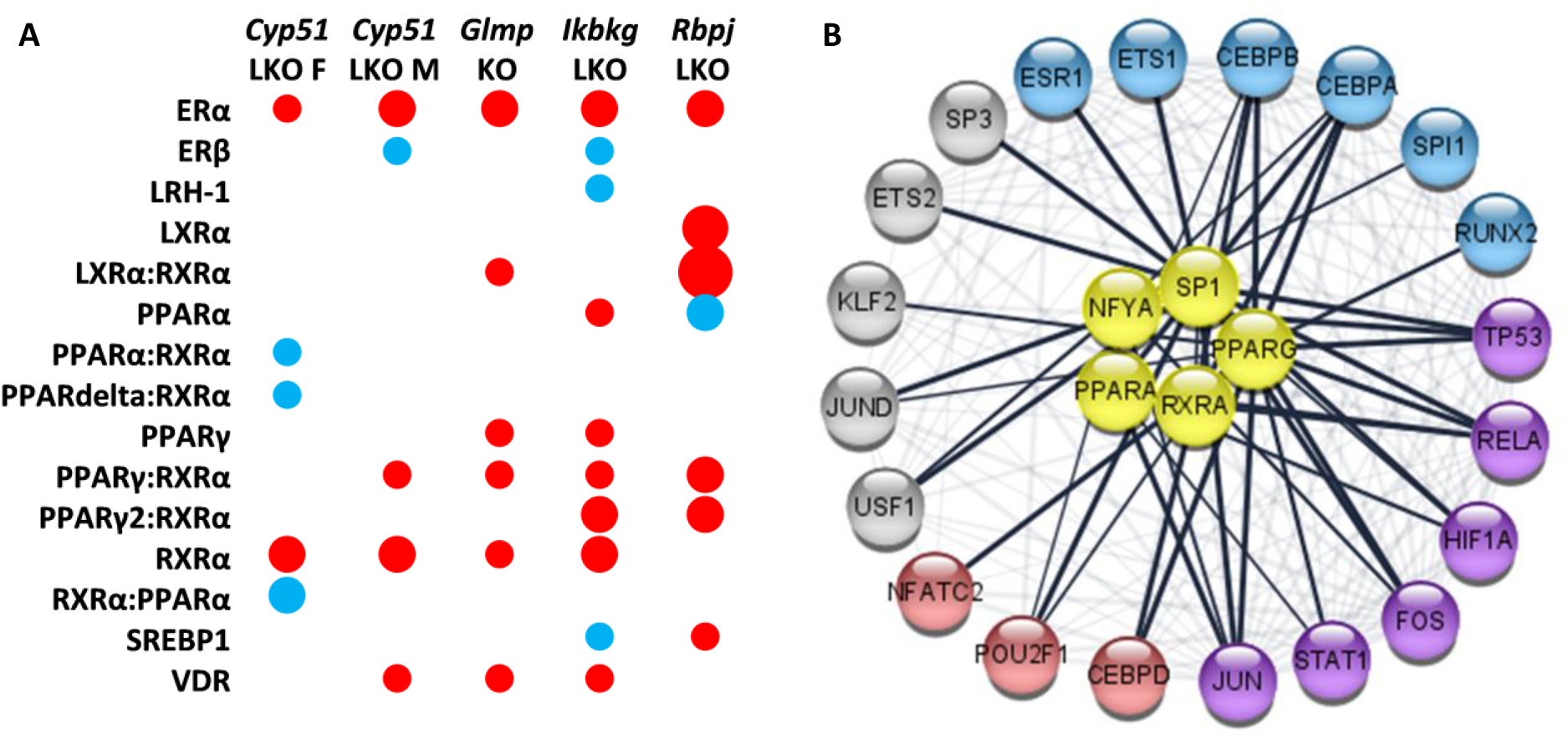
Selected enriched TFs in mouse genetic models of liver fibrosis. **A.** Model-specific enrichment of nuclear receptors in fibrotic models. Red indicates positive enrichment and blue negative. The size of the bubble reflects the fold change of each pathway. **B.** Common enriched transcription factors reveal a network-like interactions between regulators of lipid metabolism (yellow) and TFs involved in regulation of cancer pathways (blue and violet) and immune system (red and violet).

## Discussion

While it is believed that fibrosis can arise through a multitude of causes it is nevertheless reasonable to believe that a common “fibrotic programme” is hidden beneath that. To address this hypothesis, we used systems medicine approaches and pathway analyses to decipher transcriptome signatures of four genetic mouse models of liver fibrosis, one available in both sexes. In each of these models, a single gene has been knocked out and no dietary or chemical manipulation was used. Malfunction of genes with very different biological roles (cholesterol biosynthesis, Notch and NF-κB signalling, unknown lysosomal membrane protein) resulted in liver fibrosis, which progressed to liver cancer.

Herein we show that despite the different genetic insults, different sex, age, and the disease stage, a common fibrotic transcriptional programme was identified (**Fig. 5**). Positively enriched KEGG and Reactome pathways were predominantly involved in the immune system, extracellular matrix, cell-cell communication, haemostasis and cancer. This common programme is very similar to human NAFLD and NASH (Teufel et al. 2016). Downregulation of fatty acid metabolism and positive enrichment of platelets and haemostasis-related pathways is a hallmark of our data as well as transcriptomes of human NASH (Arendt et al. 2015; Mardinoglu et al. 2014; Moylan et al. 2014; Ramadori et al. 2019; Naguib et al. 2020).

**Figure 5.**
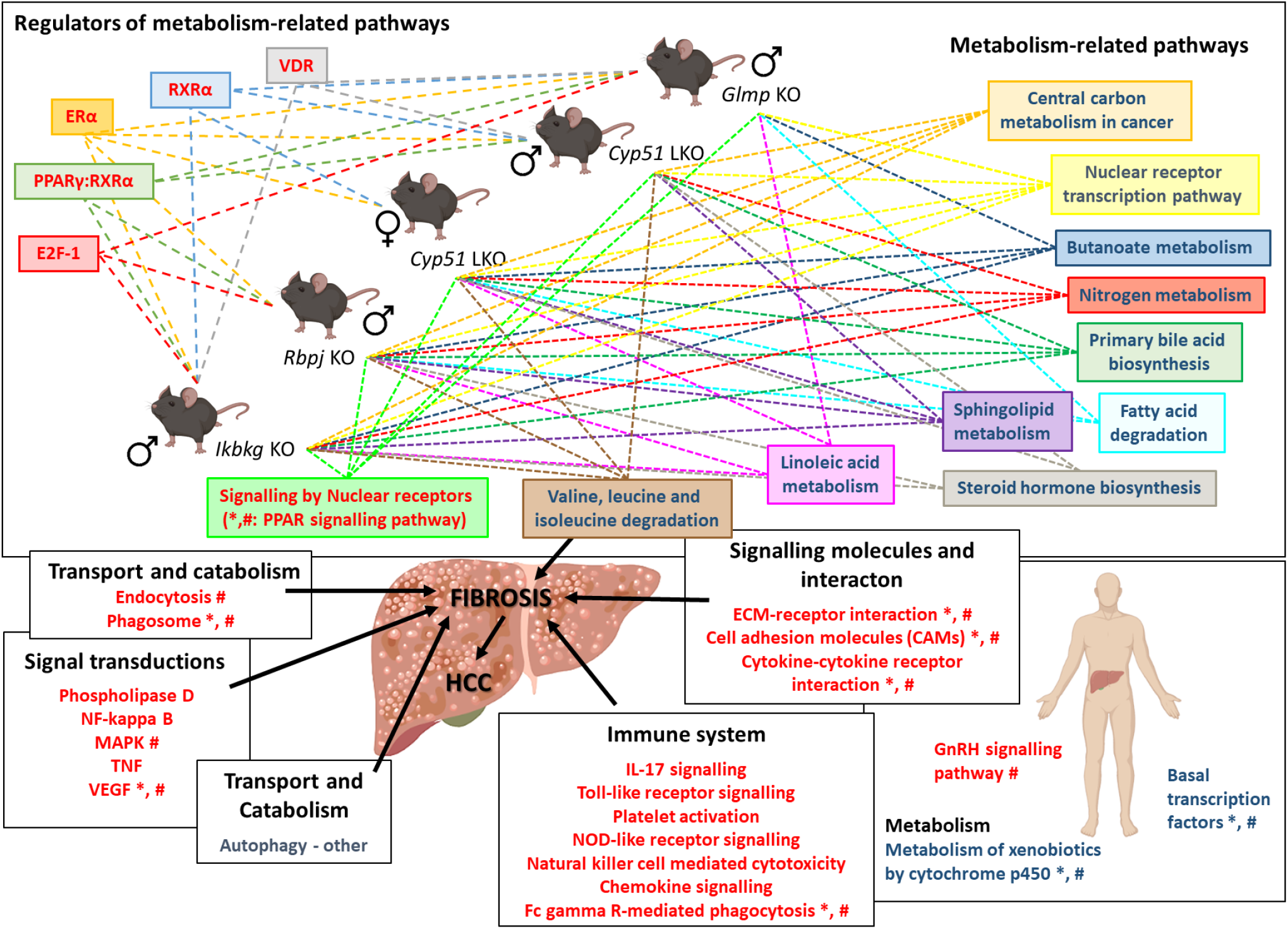
Scheme of the common transcriptional programme in mouse liver fibrotic models. Knocking out a single gene from different unrelated pathways (*Cyp51*, *Rbpj*, *Ikbkg* and *Glmp*) leads to downregulation of metabolism-related pathways, regulated by different TFs, and upregulation of signalling pathways, resulting in fibrosis. Mouse transcriptome data from mouse models was compared to human NAFLD and NASH transcriptome data (Teufel et al. 2016). * - pathways/TFs enriched also in human NAFLD; # - pathways/TFs enriched in human NASH. Blue text colour - negatively enriched KEGG/Reactome/TFs; Red text colour - positively enriched KEGG/Reactome/TFs. Hatched arrows indicate enriched pathway/TF in individual liver fibrotic mouse model. Icons are used from the BioRender library.

The negative enrichment of liver metabolic pathways indicates a molecular link between disrupted energy homeostasis and cell cycle control, which could be crucial for the development of NASH-related HCC (Satriano et al. 2019). Pathway analyses and GEMs detected wide rearrangements in the metabolism of sphingolipids, ketones, bile, linoleic and fatty acids as well as branched amino acids in all genetic models of fibrosis. These groups of metabolites present potential serum biomarkers of NAFLD/NASH progression since the changes seem to be independent of etiology of the disease. Furthermore, they represent potential new diagnostic and prognostic biomarkers of liver diseases in humans (**Table 3**). Bile acids are an example of a potential common serum biomarker of liver diseases. A potential prognostic biomarker was identified by a metabolomics prospective study where serum fatty acids, including linoleic and α-linoleic acids, were lower before the occurrence of cirrhosis in patients in comparison to healthy controls (Yoo et al. 2019). The observed model-specific changes in transcriptome signatures could reflect the unique metabolic rearrangements among fibrotic models (**Fig. 6**). For example, female *Cyp51* LKO model has upregulated genes indicating increase in GM2 ganglioside and decrease in GM3 ganglioside, while male *Cyp51* LKO, *Ikbkg* LKO and *Rbpj* LKO have potentially increased GM3 ganglioside. Overall, it seems very plausible that serum metabolites reflect the stage of liver disease and the patient’s metabolic state, and could enable differentiation of metabolic subtypes of NAFLD, NASH and beyond. For example, C4 (7-alpha-hydroxy-4-cholesten-3-one), a bile acid intermediate used to asses liver bile acid biosynthesis, was increased in obese NAFLD patients (Appleby et al. 2019), but decreased in lean patients (F. Chen et al. 2019). A combination of serum metabolites could be used for patient stratification in personalized medicine. More importantly, we need to emphasize that these genetic models develop metabolic rearrangements similar to NAFLD and NASH without obesity, dietary or chemical manipulation. We propose that overall metabolic rearrangements are crucial for the “fibrotic transcriptional programme”. However, type of the injury, stage of fibrosis and sex, define the direction, degree and type of metabolic pathway affected.

**Table 3:** Overview of changes in the level of serum metabolites and in the liver expression of transcription factors in humans with NAFLD and NASH. Selected were metabolites and TFs, which were exposed as key factors in fibrotic programme by genetic mouse models of liver fibrosis. DOWN – downregulated; UP – upregulated.

| Factor | Stage of disease | Change | Reference |
| --- | --- | --- | --- |
| **Metabolites** |  |  |  |
| Bile acids | NAFLD | DOWN | (Appleby et al. 2019; Jiao et al. 2018) |
|  | NAFLD | DOWN | (Papandreou et al. 2017) |
|  | NAFLD/NASH | DOWN | (Caussy et al. 2019) |
|  | HBV - cirrhosis | DOWN | (Arain et al. 2017) |
|  | lean NAFLD | UP | (F. Chen et al. 2019) |
| Polyunsaturated fatty acids (PUFA) | cirrhosis | DOWN | (Ristić-Medić et al. 2013) |
|  | NASH - cirrhosis/HCC | DOWN | (Yoo et al. 2019) |
| PUFA: linoleic and α-linoleic acids | NAFLD | UP | (Papandreou et al. 2017) |
|  | NASH/HCC | DOWN | (Hall et al. 2020) |
| Monounsaturated fatty acids (MUFA) | NASH - HCC | UP | (Muir et al. 2013) |
|  | lean NASH or NAFLD | UP | (Hall et al. 2020) |
| Triglycerides | lean NAFLD | UP | (Feldman et al. 2017) |
|  | lean NASH or NAFLD | UP | (Hall et al. 2020) |
|  | NASH | UP | (Alonso et al. 2017) |
| Triglycerides and cholesteryl esters | NAFLD | DOWN | (Papandreou et al. 2017) |
| Sphingolipids | NAFLD | UP | (Wasilewska et al. 2018; Apostolopoulou et al. 2018) |
|  | NAFLD/NASH | correlated with weight loss | (Promrat et al.; Ramos-Molina et al. 2019) |
|  | NASH | UP | (Alonso et al. 2017) |
| Ketones | NASH | DOWN | (Männistö et al. 2015) |
| Total cholesterol | NASH/HCC | UP | (Hall et al. 2020) |
|  | lean NAFLD | DOWN | (Feldman et al. 2017) |
| Amino acids | NASH/HCC | UP | (Hall et al. 2020) |
| Branched amino acids | NAFLD | UP | (Kaikkonen et al. 2017) |
| 6-phosphogluconic acid | NASH | DOWN | (Alonso et al. 2017) |
| **Transcription factors** |  |  |  |
| PPARα | NASH | DOWN | (Francque et al. 2015; Kersten and Stienstra 2017) |
| LXR, SREBPC1 | NAFLD | UP | (Yang, Shen, and Sun 2010) |
| RXRα | NASH | UP | (Shiota and Kanki 2013) |
|  | NAFLD/NASH | DOWN | (Allam et al. 2020) |
| PPARγ | NAFLD | UP | (Y. K. Lee et al. 2018; Westerbacka et al. 2007) |
| VDR | steatosis | UP | (Bozic et al. 2016; Barchetta et al. 2012) |
|  | NASH | DOWN |  |

**Figure 6.**
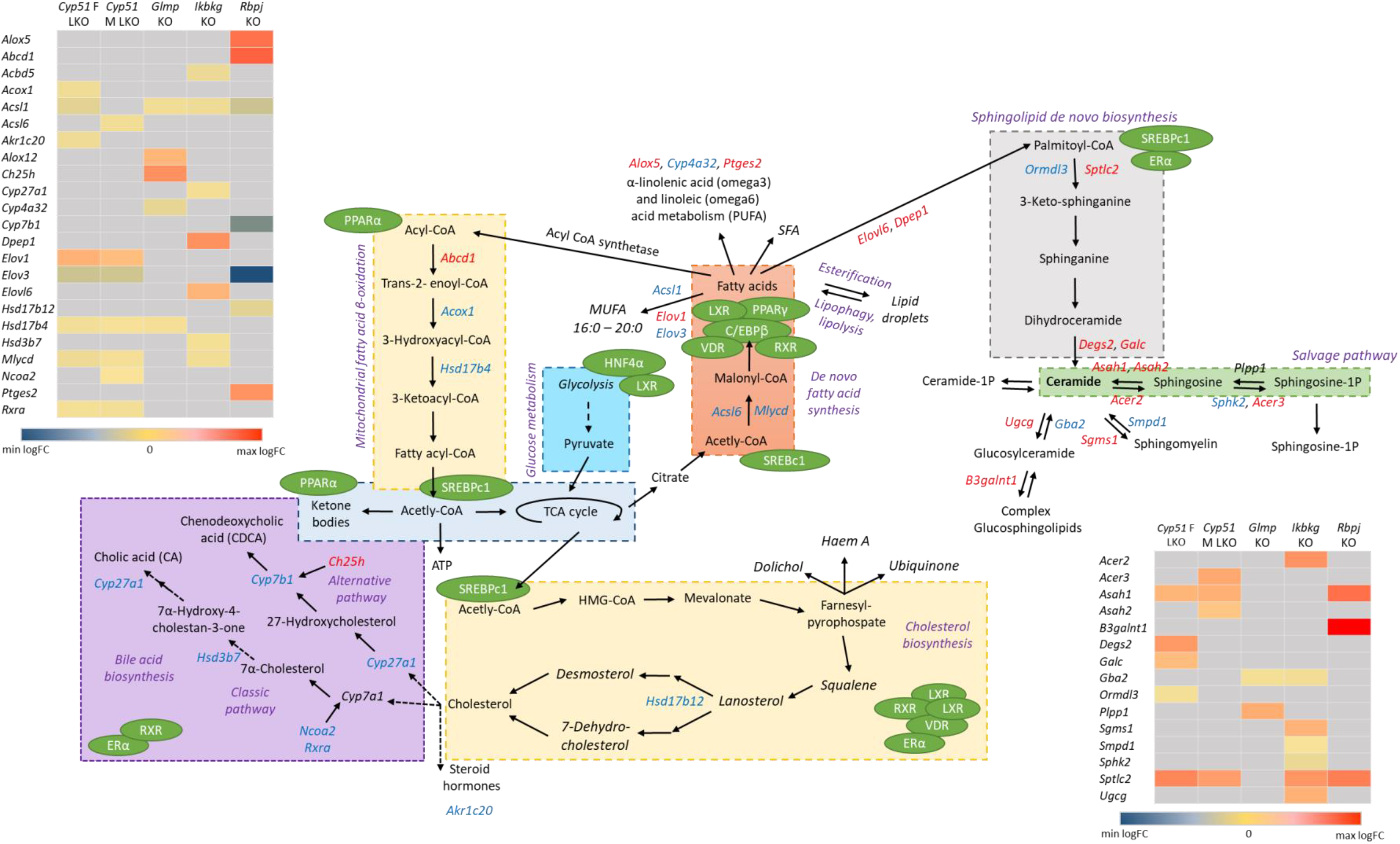
Unique lipid metabolite’s rearrangements in mouse liver fibrotic models. Pathways analyses and GEM subsystems detected model-specific deregulation of lipid metabolism in genetic models. Blue colour indicates downregulated DEG, while red are upregulated DEG. Grey in heat maps represents insignificant expression. Green circles are detected TFs in liver mouse models, which regulate different metabolic pathways (violet). Legend of heat maps are shown at bottom.

Applying different approaches and databases enabled a fresh perspective at the fibrotic transcriptome data, also in view of transcription factors as major regulators of cellular metabolism. The Reactome pathways *Nuclear receptor transcription pathway* and *Regulation of lipid metabolism by Peroxisome proliferate-activated receptor-α* were negatively enriched, while *Signalling by Nuclear receptors* and *Extra-nuclear estrogen signalling* were positively enriched. This underlines the importance of transcriptional reprogramming of metabolism in fibrosis. Recent studies exposed a suppression of liver-identity transcription factors induced by liver injury (Dubois et al. 2020). Two models, female *CYP51* LKO and *Ikbkg* LKO, exhibit a suppression of these TF. Furthermore, several TFs regulating the lipid metabolism were enriched, such as PPARγ, RXRα, VDR, PPARα, SREBF1c and LXR. Their expression is affected by NAFLD or NASH progression in human livers (**Table 3**). However, enrichment of these TF was not overlapping among genetic models, indicating that regulation of metabolism is specific to each genetic model of fibrosis and also defines the manner of metabolic rearrangements. For example, female *CYP51* LKO has the strongest inhibition of overall metabolism including TCA cycle, fatty acid and amino acid metabolism, while it is also the only model with a negative enrichment of metabolic regulators AMPK, PPARδ and PPARα. The different transcriptome landscapes resulting from different genetic backgrounds could be considered as stratified metabolic subtypes of NAFLD or NASH. This view can add in explaining the variable success of treatments targeting these metabolic regulators (Friedman et al. 2018). Based on this, we propose that regulators and their downstream metabolites that differ between the genetic models warrant further testing as potential biomarkers in a human setting, to enable stratification of patients for metabolic subtypes of fibrosis. This could substantially increase the impact of existing and novel therapeutic strategies.

Epidemiological data in humans clearly shows that estrogen has a protective role against NAFLD and NASH in premenopausal females (C. Lee, Kim, and Jung 2019). Thus, sex hormones affect the development of liver diseases (Ruggieri, Gagliardi, and Anticoli 2018). Our pathway and TF enrichment analyses and GEMs exposed changes in overall steroid hormone metabolism as one of the hallmarks of the fibrotic programme. *Extra-nuclear estrogen signalling* and ERα were positively enriched in all mouse models, regardless of sex, while *Steroid hormone biosynthesis* was negatively enriched in four models. Data in humans confirm increased expression of ERα in NAFLD livers and its correlation with the severity of steatosis (Choi et al. 2018). ERα knockout in mice present with induced steatosis in both sexes, indicating that ERα activation in fibrosis could be a sex-independent protective adaptation against liver insults, exposing estrogen receptor as a potential drug target for NAFLD management (Hart-Unger et al. 2017; Qiu et al. 2017; Hevener, Clegg, and Mauvais-Jarvis 2015). Interestingly, even though we have previously shown a sex-dependent difference in the progression of liver fibrosis (Urlep et al. 2017), we observe similar changes in steroid-related pathways in both sexes (Lorbek et al. 2015). Based on our analysis we cannot make conclusions regarding fibrotic programmes and each sex since too little data is available for the females. Signalling through the estrogen receptor seems to be a part of the common fibrotic transcriptional programme, regardless of the insult or sex. It is necessary to have more data available for both sexes. An important part of stratification would be if one could prove that the liver fibrotic transcriptome signatures differ between females and males.

Since one of the fibrotic models has a knockout in the gene *Glmp* with not fully understood function, the comparative transcriptome analysis helped in shedding light at its role at the molecular level. The full knockout of GLMP leads to liver fibrosis with inflammation, oval cell activation and proliferation, hepatocyte apoptosis, oxidative stress and development of HCC and hemangioma-like tumours from the age of 12 months (Nesset et al. 2016; Kong et al. 2014). Since evidence for HCC was not substantiated in KEGG analyses, in contrast to other mouse models, we anticipate that alternative pathways are responsible. A likely explanation for liver cancer in *Glmp* KO mice is impaired autophagy due to deficiency of lysosomes which may on one hand contribute to the pathogenesis of NAFLD (Ke 2019), and can also lead to lysosomal storage disorders (Seranova et al. 2017). A recent report demonstrated that MFSD1 and GLMP, both lysosomal membrane proteins, interact and affect the expression of each other (López et al. 2019). MFSD1 belongs to a group of proteins transporting nutrients, waste and ions across membranes (Perland et al. 2017). A dysfunctional GLMP/MFSD1 complex could induce abnormal functions in lysosomes, severely affecting autophagy (Yim and Mizushima 2020). We propose that disturbances in these two pathways work cooperatively in increasing the ER stress, inflammatory-related KEGG pathway and development of fibrosis (Lavallard and Gual 2014). This is corroborated by the observed negative enrichment of mTOR signalling and KEGG pathways associated with detoxification, such as cytochromes P450.

**In conclusion**, based on comparing different genetic models of liver fibrosis without dietary manipulation, we revealed common liver fibrotic transcriptome signatures with high similarity to signatures of human NAFLD and NASH. A hallmark of the fibrotic programme are changes in metabolic pathways related to lipids, such as bile acids, steroids, sphingolipids and fatty acids. These metabolites and their regulators (AMPK, FOXOA1, SREBP1, LXRα, PPARδ and PPARα) exhibit enrichment that depends on the genetic background, exposing their potential to serve as diagnostic and prognostic biomarkers of fibrotic subtypes also in humans. They could also enable a more precise metabolism-related stratification of NAFLD/NASH patients before entering clinical drug trials and facilitate the implementation of personalized liver disease management.

## Materials and Methods

### Microarray-based gene expression analysis

*Glmp* KO and *Cyp51* LKO transcriptome were assessed in-house, while raw transcriptome data for *Ikbkg* LKO and, *Rbpj* LKO models were obtained from GEO. To assess the *Glmp* transcriptome, we hybridized Affymetrix GeneChip Mouse Gene 2.0 ST Arrays (Affymetrix, Santa Clara, California, USA) with samples from livers of 16 *Glmp KO* and *Glmp WT* mice at age 8 and 18 weeks. Each group consisted of 4 samples (**Suppl. Table 8**). Data analysis were performed using R and Bioconductor software packages. We normalized raw (CEL) expression data using the RMA algorithm from package *oligo* (Carvalho and Irizarry 2010). Quality control and outlier detection were performed using package *arrayQualityMetrics* before and after normalization (Kauffmann, Gentleman, and Huber 2009). Raw as well as normalized data were deposited to GEO under accession number GSE154021.

The generation of transcriptome data from 19-week *Cyp51* LKO (GSE58271), 4-week *Rbpj* LKO (GSE121302), 8) to 9-week *Ikbkg* LKO (GSE33161) and mice were described before (Lorbek et al. 2015; Tharehalli et al. 2018; Cubero et al. 2013). RMA algorithm from package *oligo* and quantile normalization from package *limma* (Smyth 2004) were used for re-normalization of raw expression data from Affymetrix and Agilent arrays, respectively. Package *limma* was used to fit individual normalized gene expression data using linear regression models as shown in **Suppl. Table 8**. Empirical Bayes statistics were used to estimate the statistical significance of expression differences of genes and Benjamini-Hochberg procedure was used to control false discovery rate (FDR) of differential expression at α=0.05.

KEGG pathways (Kanehisa et al. 2019), Reactome pathways (Jassal et al. 2020) and TRANSFAC database ver. 2020.1 (Matys 2006) were used for functional enrichment studies. Gene sets containing 5 or more elements were constructed and tested for enrichment using the PGSEA package (Furge and K, n.d.). In the case of transcription factor (TFs) enrichment, factors were merged based on their ID irrespective of their binding sites. Statistical significance of gene set enrichment was estimated using the same approach as for individual genes.

To facilitate comparative functional genomics analysis differentially expressed genes (DEG) and enriched gene sets were partitioned according to their overlaps between the studies. Genes and gene sets were split to up/down regulated and positively/negatively enriched and their numbers are reported. Overlaps between the models are visualized by Venn using package *VennDiagram* (H. Chen 2018).

Functional similarity between the mouse models was quantified as a ratio of significant expression/enrichment changes that are in common to the models vs. the significant expression/enrichment changes of each model individually, e.g., the number of DEG in the intersection of two models was divided by the number of DEG for each model. Thus, a non-symmetric similarity matrix was calculated summarizing similarities between all pairs of models from a perspective of each model. Ratios of significant DEG are shown in **Suppl. Table 3** (non-diagonal values marked with green), together with the number of DEG (diagonal values that are >1 and marked with red). Furthermore, the similarity between models is expressed separately for positive and negative expression/enrichment changes; thus each pair of the model is characterized by two ratios, left for positive and right for negative expression changes. Ratios are represented row-wise: the number of DEG in common was divided by the number of DEG of the model within the corresponding row. **Suppl. Table 3** show similarities between the models for KEGG and Reactome pathways and TFs, respectively.

### Genome-scale metabolic modelling

We performed the integration of DEG into the genome-scale metabolic model (GEM) of C57BL6/J mice liver tissue, which was previously described (Mardinoglu et al. 2014) and is available in the Metabolic Atlas Database (www.metabolicatlas.org) (Pornputtapong, Nookaew, and Nielsen 2015). DEG were integrated into the model using the Metabolic Adjustment by Differential Expression (MADE) method (Jensen and Papin 2011; Jensen, Lutz, and Papin 2011). MADE integrates differential expression data into a reference model using the flux balance analysis (FBA) to obtain a functional metabolic model describing a perturbed state of a system (e.g., after gene silencing). When reference and perturbed models are available, up-/down-regulated reactions can be identified using the flux variability analysis (FVA) (Gudmundsson and Thiele 2010). The list of up-/down-regulated reactions obtained with the FVA was used to perform the metabolic subsystem enrichment analysis based on the hypergeometric test. The Benjamini and Hochberg procedure was used for the P values adjustment. The cut-off value for significantly up-/down-regulated subsystems was set to 0.05.

## Acknowledgements

We would like to acknowledge Nejc Nadižar for the help with generating figures.

## Competing interest

The authors declare no competing financial or non-financial interests or conflicts of interest.

## Notes

### Competing Interest Statement

The authors have declared no competing interest.

